# Eco-evolutionary Guided Pathomics Analysis to Predict DCIS Upstaging

**DOI:** 10.1101/2024.06.23.600274

**Authors:** Yujie Xiao, Manal Elmasry, Ji Dong K. Bai, Andrew Chen, Yuzhu Chen, Brooke Jackson, Joseph O. Johnson, Robert J. Gillies, Prateek Prasanna, Chao Chen, Mehdi Damaghi

## Abstract

Cancers evolve in a dynamic ecosystem. Thus, characterizing cancer’s ecological dynamics is crucial to understanding cancer evolution and can lead to discovering novel biomarkers to predict disease progression. Ductal carcinoma in situ (DCIS) is an early-stage breast cancer characterized by abnormal epithelial cell growth confined within the milk ducts. In this study, we show that ecological analysis of hypoxia and acidosis biomarkers can significantly improve prediction of DCIS upstaging. First, we developed a novel eco-evolutionary designed approach to define habitats in the tumor intraductal microenvironment based on oxygen diffusion distance. Then, we identified cancer cells with metabolic phenotypes attributed to their habitats, including CA9 for hypoxia responding phenotype, and LAMP2b for acid adapted phenotype. Traditionally these markers have shown limited predictive capabilities for DCIS progression, if any. However, when analyzed from an ecological perspective, their power to differentiate between non-upstaged and upstaged DCIS increased significantly. Second, we discovered distinct niches with spatial patterns of these biomarkers and used the distribution of such niches to predict patient upstaging. The niches were characterized by pattern analysis of both cellular and spatial features. With a 5-fold validation on the biopsy cohort, we trained a random forest classifier to achieve the area under curve (AUC) of 0.74. Our results affirm the importance of tumor ecological features in eco-evolutionary-designed approaches for novel biomarkers discovery.

**Significance:** Our results affirm the importance of spatial and ecological features in eco-evolutionary-designed biomarkers discovery studies in the era of digital pathology.

## Introduction

In recent years, the understanding that cancer is a dynamic ecological and evolutionary process has become deeply entrenched (1,2,3). To date, several evolutionary approaches have been adapted and applied in cancer biology; however, tumor ecosystem and ecological studies are still overlooked (3,4). Within the human body and much like organisms in the nature, cancer cells follow ecological principles, utilizing resources and establishing niches within tissues habitats (5,6). This ecological perspective of cancer is crucial for discovering the natural selection driving cancer evolution. Such insights may potentially lead to improved cancer prognosis, progression prediction, risk stratification, and therapeutic strategies. If tumor evolutionary and/or its ecological state could be reliably achieved using a single biopsy tissue, clinical translation would be comparatively more manageable. Nevertheless, studies have yet to determine whether measures of tumor eco-evolution derived from a single biopsy sample are adequate, or if the inclusion of multiple samples significantly enhances predictions of clinical outcomes (7).

Breast cancer incidence in the US has been increasing over the past decade at a rate of 0.5% per year (8). With increased mammographic screening, there has been a substantial increase in detecting the early non-invasive forms of breast cancer, such as ductal carcinoma in situ (DCIS)(2,9). About one-third of breast cancers detected by mammography are DCIS (10). DCIS and IDC (invasive ductal carcinoma) are indistinguishable by (epi-)genetic mutations, gene expression, or protein biomarkers. It is also not possible to predict whether DCIS will remain non-upstaged or progress to aggressive disease. Therefore, almost all early tumors are treated the same (2,12–14). To avoid such overtreatment for non-upstaged group, more research is needed to fully understand evolution from pre-cancer to non-upstaged DCIS or progress to IDC (9).

DCIS can be defined as a heterogeneous group of neoplastic lesions confined to the mammary ducts. The confinement of proliferating neoplastic cells inside the duct and growth of pre-cancer cells toward the center of the duct, which is far from vasculature, causes limitations in oxygen access. This intraductal oxygen availability is also influenced by complex ecosystems surrounding the duct, such as vascular density and activity (15), stiffness of extracellular matrix (ECM) (16), and metabolites (6,17,18,19) (**Figure 1A**). Local microinvasion is the main difference between DCIS and IDC and might also be the first evolutionary step of progressing in the case of linear evolution(11). Microinvasion consists of cohorts of cancer cells that breach the basement membrane into the surrounding ECM (6,11,18,20). We hypothesize that non-genetic ecological factors, such as intra-ductal microenvironment, may be responsible for transitioning from DCIS to IDC. To validate this hypothesis, we propose a novel method to study DCIS evolution, by capturing and characterizing tumor ecological features such as “habitats” and “niches” in breast ecosystem (21). We started by defining the habitats based on availability of oxygen into: a) oxygenated habitat and b) hypoxic habitat defined by distance from the duct boundary (18, 22). We then defined local niches using biomarkers indicative of cancer cell response to variation in oxygen availability or acid adaptation. These biomarkers are designed based on our prior cancer eco-evolutionary studies and findings. Oxygen availability determines the source of energy production as of either mitochondrial respiration or glycolysis. Hypoxic cells switch to glycolysis, causing lactic acid production that can lead to acidosis when lactic acid is locally accumulated. Peri-luminal cells will experience hypoxia if they are far (>0.125 - 0.160 mm) from a blood supply. These cancer cells inhabit a microenvironment of hypoxia, acidosis, and severe nutrient deprivation (18,22). Our previous studies strengthen the acid-induced evolution model of breast cancer and our proposed evolutionary designed biomarkers including CA9 and LAMP2b in this translational research (6,17,20,23,24). Here we examined the role of these biomarkers within an eco-evolutionary concept as a predictor of DCIS upstaging for the first time. We used these markers as representative of the cancer cells metabolic states to define niches that can select for more aggressive phenotypes, leading to microinvasion and DCIS upstaging to IDC. Niches were defined as clusters of cells with similar phenotype responding to hypoxia and/or acid.

**Figure 1.**
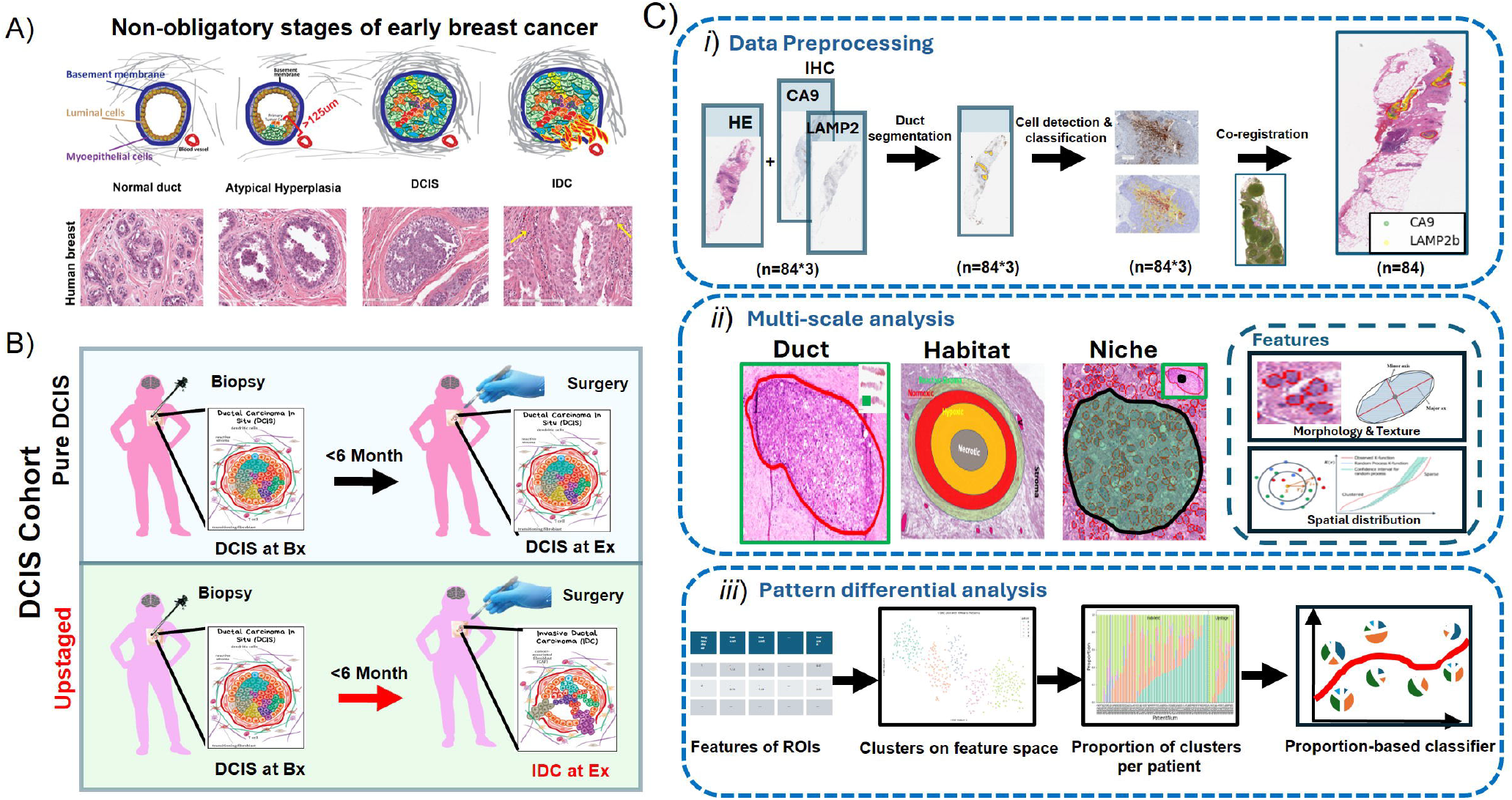
Ecological and evolutionary designed biomarkers of DCIS upstaging. **A)** Model of microenvironment-driven evolution of breast cancer from normal breast tissue to DCIS and IDC: Our schematic is overlaid on HE staining of breast cancer specimens at different stages of DCIS and IDC. Different patients may experience various types of evolutionary trajectory following different evolutionary models, including linear and branched progression from DCIS to IDC shown here. Note that these events are not sequential or stepwise. **B)** The patient cohort was curated from retrospective DCIS samples, with two sample collections at biopsy and excision. The main criterion was the diagnosis of DCIS at the biopsy stage. **C)** Eco-evolutionary designed-machine learning assisted pipeline to define cancer cell niches inside oxygen habitats in DCIS. ***i*)** Data preprocessing steps including duct annotation, cell detection and classification for HE and IHC slides, followed by co-registration to map IHC-identified cells onto the HE slides. ***ii*)**The analysis is carried out at multiple scales, namely duct, habitat and niche, from the largest to smallest. At each scale the nucleus morphology texture feature and spatial features are extracted. ***iii*)** The pattern differential analysis approach where the patterns are firstly identified and then the proportions of such patterns are used as features to predict the upstaging status of a patient.

To perform our analysis, we curated a retrospective cohort of DCIS patients, with specimens collected from Biopsy (Bx) samples before surgery and the whole tumor after Excision (Ex). All the patients had histologically confirmed DCIS at core Bx, followed by diagnosis confirmed on surgical excision specimens with either DCIS or IDC (**Figure 1B**). Our prediction model is trained and tested on the Bx samples. We first stained 3 sequentially sections for hematoxylin and eosin (HE), CA9 ab, and LAMP2b ab. We then manually annotated every single duct bigger than 400 μms in diameter, 200 μms in radius. The 200 μms in radius annotation ensures each duct has both oxygenated and hypoxic habitats to build a balanced cohort for analysis. Then we scored the positivity of each habitat for both CA9 and LAMP2b to compare both non-upstaged and upstaged group. We finally developed a novel algorithm to detect intra-ductal niches based on biomarker expression similarity. We applied multiple spatial functions and spatial entropies to define niche and micro-niches describing the spatial patterns of the positive cells. Thus, we studied the spatial organization of CA9- and LAMP2b-positive cells at four different scales: whole slide, duct, habitats, and niches. After a systematic and comprehensive analysis, we observed that the spatial features at the finest niche level possess the most predictive power where the micro-niches were defined by the expression of both CA9 and LAMP2b. By characterizing these niches and micro-niches with spatial and pathomics features, we then developed a risk scoring system by integrating pathological imaging and molecular features of early-stage breast tumors (**Figure 1C**). We show that quantitative analyses of immuno-histological images combined with the tumor’s eco-evolution dynamics and underlying molecular pathophysiology can significantly improve predicting if the neoplasm is invasive disease. Our developed machine learning model, fine-tuning the tumor ecosystem into habitats and niches, provides informed decision support.

In summary, we show that eco-evolutionary applications can improve existing models of DCIS progression. Our developed machine learning techniques to calculate the niches and habitats inside the tumor of DCIS patients with pure DCIS and upstaged disease can provide supportive information to understand the disease progression. Finally, we suggest that deploying eco-evolutionary principles and machine learning techniques, can propose a novel consilient approach to improve stratifying DCIS patients.

## Materials and Methods

### Method overview

Our evolutionary analysis pipeline takes 3 sequential sections of each patient sample, defines oxygen habitats (normoxia and hypoxia) and abundance of CA9 and LAMP2b cells positives in those habitats, detects intra-ductal cell niches, characterizes these niches’ spatial and morphological features, and finally use these eco-evolutionary markers to predict whether the patient will be upstaged or not. In particular, the pipeline has 4 modules. First, we annotate single ducts to their habitats and score habitats. Then align ducts from all three whole slide images (WSIs). This ensures cells of different slides are aligned and we can characterize their interactions in habitats and niches. In the second module, we detect and map all eco-evo positive cells (i.e., cells positive for CA9 and LAMP2b) into the same duct on HE slide and detect different clusters of cells as niches. In the third module, we characterize these niches with comprehensive spatial statistical features, as well as their morphological features as observed in HE. Finally, we categorize these niches into different subclasses through deep learning-based dimension reduction and feature-based clustering. We also use the distribution of niche subclasses to characterize different samples/patients. We demonstrate the discriminative power of these niche-based characterization in predicting whether a patient will be upstaged or not. **Figure 1C** illustrates the overview of our pipeline.

### Data preparation and usage

Data used in this study comes from biopsy samples collected after mammography and before surgery. 84 samples including 68 pure DCIS and 16 progressed to IDC were collected. This study complied with the Health Insurance Portability and Accountability Act and was approved by the institutional review board, with a waiver of the requirement for informed consent. Women with a core biopsy diagnosis of DCIS between 2012 and 2022 who consented to Moffitt Cancer Center Total Cancer Care protocol were included in this analysis. Cases were excluded if surgical excision was performed more than 6 months after the core biopsy, if there was concurrent ipsilateral invasive breast cancer or metastatic malignancy, or if neoadjuvant chemotherapy (for a concurrent contralateral breast malignancy) or chemotherapy for any malignancy was administered between the dates of the DCIS core Bx and surgery. We also excluded the patient if there was a personal history of IDC or DCIS within 12 months prior the Bx or a concurrent diagnosis of Paget disease in the ipsilateral breast. After applying these inclusion and exclusion criteria, 84 cases of biopsy-proven DCIS were identified, of which 16 were upstaged at surgery and 68 remained non-upstaged. From each patient one block selected with at least one lesion in it. From each block, we obtained 3 whole slide images, including 1 HE and 2 IHC slides. To minimize confounding effects, non-upstaged and upstaged patients were matched across clinical features including age, race, ethnicity, tumor grade, and ER/PR status. Demographic characteristics of the cohort (e.g., age range, race/ethnicity distribution) are summarized in **Supplementary Figure S1**.

### Sample selection, immunohistochemistry and HE staining

Tumor blocks were selected by pathologists using the archived HE stained slides. The blocks were sequentially sectioned 4 μm and deidentified for research use. 3 slides were stained with primary antibodies of 1:100 dilution of anti-LAMP2 (Abcam Cat# ab18529, RRID:AB_2134632)), and 1 ug/ml concentration of anti-CA9 (R&D Systems Cat# AF2188, RRID:AB_416562), and HE staining using standard hematoxylin and eosin protocol. Normal placenta was used as a positive control for LAMP2b and clear cell renal cell carcinoma was used as a positive control for CA9. For the negative control, an adjacent section of the same tissue was stained without application of primary antibody and any stain pattern observed was considered as non-specific binding of the secondary. Primary immunohistochemical analysis was conducted using digitally scanned slides. The scoring method used by our pathologist to determine (a) the degree of positivity scored for each sample ranged from 0 to 3 and was derived from the product of staining intensity (0 – 3+). A zero score was considered negative, score 1 was weak positive, score 2 was moderate positive, and score 3 was strong positive. (b) The percentage of positive cells (on a scale of 0-3).

Whole slide imaging (WSI) of IHC and HE slides were obtained by scanning at 20X magnification (of 0.5022 micrometer per pixel) using Aperio AT2 from Leica Biosystems. Images were transferred to cloud storage and locally to be uploaded in QuPath (v0.4.3; RRID:SCR_018257) software for analysis. QuPath software was used to detect the positive pixels for each IHC marker (CA9 and LAMP2b) and to segment the HE images into hypoxic and normoxic tumor habitats based on their distance from the basement membrane. The ‘Positive Cell Detection’ function from QuPath was used to automatically classify the positivity of CA9 and LAMP2b markers and validated by the study pathologist.

### MODULE 1: Duct annotation and alignment

#### Manual annotation of ducts

We annotate and align ducts within all input slides (1 HE + 2 IHCs per sample). After annotating ducts from WSIs of all three modalities, we align and co-register the ducts from the three modalities. This alignment enables us to map cells into the same spatial domain in each slide and analyze their interaction. QuPath was used as the interface to annotate ducts by the pathologist (Dr. Bai) and reviewed by Dr. Damaghi. To ensure the same number of oxygenated and hypoxic habitats, we only select ducts of >400 μm diameter from basement membrane. Following this, based on distance, each duct was annotated with four layers: adjacent stroma, oxidative/normoxia, hypoxic/hypoxia, and necrosis. Within the duct, necrosis was defined as any area containing dead cells, as identified by a lack of nuclei. Oxidative layer was defined as the area containing cells inside the duct within 125 μm of the basement membrane. Hypoxia was defined as the area containing cells inside the duct further than 125 μm from the basement membrane. The annotations were done for all 84 samples in the Bx cohort, and then were exported as standard GeoJSON files. Pathologists were blinded to patient outcomes during annotation and scoring.

#### Co-registration

To build a multiplex image from monoplex slides, we register both CA9 and LAMP2b IHC slides towards the HE slides. For that reason, we applied direct co-registration at the whole slide level with manual landmarks for alignment at each duct. We further co-register the slides in a duct-by-duct fashion. Using initially registered whole slides, and spatial proximity, we identify the corresponding ducts at the HE and 2 IHC slides. Next, we register both the CA9 duct and LAMP2b duct into the corresponding HE ducts. We use Virtual Alignment of pathology Image Series (VALIS), which provides a fully automated pipeline to register whole slide images (WSI) using rigid and/or non-rigid transformations (34). For each sample, we chose non-rigid registration and registered the ducts from CA9 and LAMP2b towards the reference HE ducts. The co-registration procedure and the qualitative results are shown in **Figure S4**. The co-registration provides a mapping of any cells detected in CA9 or LAMP2b towards a shared spatial domain, enabling the analysis of their interactions.

### MODULE 2: Cell detection and niche definition

#### Cell detection

With the duct annotations in place, we automatically detect cells from the 2 IHCs and determine if they are positive in CA9 or LAMP2b based on their intensities. For each duct, we detect cells using QuPath watershed cell detection algorithm (25). Based on the intensity level, we categorize the cells into 4 groups: ‘Negative’, ‘1+’, ‘2+’, and ‘3+’. The detection of cells within a duct is done by StarDist (25,35) extension in QuPath.

#### Graph construction for niche detection

After annotating all the positive cells (i.e., CA9 or LAMP2b positive cells), they were mapped on HE slides, enabling us to detect niches on HE slides. We construct a graph with the positive cells by connecting cells whose distances are smaller than a certain threshold and detect connected components of the graph as representatives of cells living in “niches”. Multiple thresholds have been experimented and an optimum value is selected based on performance. Each niche was set to have cells with similar eco-evo phenotype and be spatially coherent. Therefore, we overlay both CA9 and LAMP2b positive cells into the same domain as an approximation of the local eco-evo cell distribution (**Figure S5**). This gives us the opportunity to measure their interaction via spatial statistical functions as defined later. Based on the same principle, we use cell morphological features extracted from HE within the region of each niche to characterize the niche.

### MODULE 3: feature extraction from niches for characterization

Once niches are detected, we extracted both spatial and morphological features. Then we characterize spatial interaction patterns, utilizing various spatial functions as features. This is extra to the cell features consisting of morphology and texture features that adopted in HE images.

For cellular features we measured both morphological and texture features. The morphological features include area, eccentricities, circularity, elongation, extent, major axis length, minor axis length, solidity and curvature. The texture features include angular second moment (ASM) of co-occurrence matrix, contrast, correlation, entropy, homogeneity and intensity. All features were calculated following the implementations in the sc-MTOP package (36).

Although we do not have exact cell-to-cell correspondence between the cells within a niche and cells detected in HE, we still can aggregate morphological and texture features within the proxy of the cells part of a niche to characterize that niche. For each niche, we identify the concave hull region enclosing its eco-evo positive cells within a duct on HE slide. Next, we aggregate cell features across all HE-detected cells within the corresponding region. For each cell feature dimension, we calculated its mean, standard deviation, maximum, minimum, kurtosis and skewness.

#### Spatial features

We extract various spatial statistical functions (37) to characterize residing cells and their interactions to define niches. These functions are listed below:

##### G Function

The G function, denoted as G(r), is the cumulative distribution function of nearest-neighbor distance. The G function provides insights into the clustering or dispersion behavior of the point pattern.

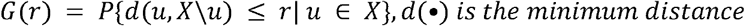

##### F Function

The F function, known as the empty space function, is the cumulative distribution function of the empty-space distance. The F function is commonly used to assess the regularity or inhibition patterns in point patterns.

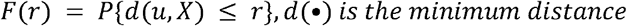

##### K Function

Ripley’s K function, denoted as K(r), is a measure of second-order intensity or spatial interaction. It assesses whether points tend to be more clustered or dispersed within a certain distance r compared to a CSR process. It considers both the distance and intensity of points to capture the clustering behavior of the point pattern.

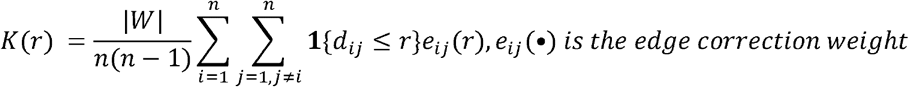

##### L Function

L function is a variance stabled version of K function.

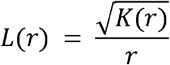

We calculated G, F, and L functions in both univariate and multivariate fashions. For each of the functions, the distances between source cell and the target cells are considered. Univariate spatial functions sample source cells and target cells from the same type of cells while multivariate counterparts’ sample from different types of cells. Univariate G, F, and L are calculated for the single-marker cell subsets, and multivariate G_cross and L_cross for different subsets such as CA9-LAMP2b. ‘Gest’ function and ‘Fest’ function from ‘spatstat’ R package (v3.3-1; https://CRAN.R-project.org/package=spatstat) were used with Kaplan-Meier estimator (38), and ‘Lest’ function was used with isotropic correction (39, 40).

### MODULE 4: Diagnostic risk estimation with pattern proportion

We trained a classifier using the niches defined and characterized as described above to predict whether a patient will be “upstaged” or “non-upstaged”. This establishes the diagnostic power of the identified niches. Worth to note that a direct aggregation of niche information within each sample/patient is not sufficient. One technical challenge is that the niche features computed in the previous module are high dimensional and the niche features are diversely distributed. To solve that we first find a simplified distributional description of the niches, and then use the simplified description for prediction. First, we cluster the niches into different sub-classes based on their features. The clustering is carried out using K-means clustering with a tunable parameter k. Once the niche sub-classes are determined, we use their distribution on each sample to predict its upstaged/not-upstaged status. The prediction power of the classifier sheds light on the diagnostic power of the niches and their spatial and cellular features. Five-fold cross-validation was employed, with one-fold designated as the test set in each run. This approach prevents data leakage and helps mitigate overfitting. To understand the contribution of each feature to the prediction model, we employed SHAP (SHapley Additive exPlanations) analysis. Machine learning models were implemented using scikit-learn (RRID:SCR_019053)

## Data and Code Availability

The data generated in this study are available within the article and its supplementary data files. All the images and their annotations at duct or habitat levels are deposited in the physical sciences in oncology network (PSON) and can be accessed at DOI: https://doi.org/10.1016/j.isci.2024.109433. Custom Python and R analysis code is available at https://github.com/sbu-damaghi-ceel/DCIS_archive.

## Study Limitation

This study excludes unaccounted clinical variables that can play role in DCIS upstaging. Another limitation is the absence of independent external validation due to our particular annotations.

## Results

### Sample curation and cohort building

We built a retrospective cohort of 84 patients with histologically confirmed DCIS on core Bx, followed by surgical Ex, with available FFPE from Moffitt Cancer Center Biobank. The cohort has two arms: i) pure DCIS: the patient diagnosed with DCIS at both Bx and Ex; ii) upstaged: patient with DCIS at Bx and IDC at Ex (**Figure 1B**). HE slides of Bx cores were retrieved from the biobank at the Moffitt

Cancer Center tissue core and reviewed by a study pathologist (43). Then the selected blocks were pulled and sequentially cut for HE, CA9, and LAMP2b staining. The HE and subsequent 2 IHC slides are digitally scanned using the Aperio XT® high-throughput slide scanner and housed on the web-based Aperio server/Spectrum database package. Upstage status was pulled from the electronic medical record and confirmed by our study pathologist from the Ex tissues (**Figure 1C**). All images were then segmented and annotated using QuPath supervised by study pathologist (25) (43).

Non-upstaged and upstaged patients were matched across clinical features, including age, race, ethnicity, grade, and ER/PR status, to minimize their influence on the analysis (**Figure S1**). To validate the comparability of these groups, we conducted a Wilcoxon rank-sum test for the continuous variable (age) and Chi-square tests for the categorical variables (race, ethnicity, grade, ER/PR status). None of these tests showed significant differences between the two groups, with all p-values larger than 0.1, indicating that the groups were well-matched (**Figure S1**).

### Annotation and eco-evolutionarily mapping of habitats at the individual duct level

We have shown previously that peri-luminal cells that are far (>0.125 - 0.160 mm) from a blood supply inhabit a microenvironment of hypoxia and acidosis (18,20,26). Thus, we created two simple annotation zones on HE slides based on oxygen diffusion distance representing oxygen habitats: i) hypoxic zone or habitat that is above 125 μm from the duct boundary, basement membrane, and ii) normoxic habitat that is the outer regions adjacent to the basement membrane (**Figure 2A**). We used the basement membrane as our zero point of reference. We also annotated necrotic zones inside the hypoxic habitats that also represent the anoxic habitat falling perfectly above 0.160 mm distance from basement membrane. Since adjacent stroma is also of interest to our group and others, we annotated adjacent stroma for each duct with binary scoring of 1 for having adjacent stroma or 0 for lacking it (**Supplementary Table 1**). To ensure a balanced representation of hypoxic and normoxic habitats, we excluded small ducts by establishing a duct size threshold of minimum 400 μm in diameter (or 200 μm radius) for manual annotation (**Figure S2**). After annotating all the ducts bigger than 200 μm of radius on HE slides, we expanded our annotations to other 2 consecutive IHC slides stained with CA9 and LAMP2b antibodies (**Figure 2B**). Subsequently, our pathologist, Dr. Bai, manually scored each duct for their architecture (Solid form, Cribriform, Papillary, and Comedo), nuclear grade, necrosis, microcalcification, reactive stroma, and lymphocyte aggregate. Interestingly inside one patient we found heterogeneity of ducts as they are different tumors implying the possible multiclonality. She then scored hypoxic and normoxic habitats based on CA9 and LAMP2b positivity using a scoring scale of 0–3 (**Supplementary Table 1**). Following this, positive cells in IHC slides were counted using QuPath (25), habitats were categorized into different classes based on the count of positive cells. The distribution of these habitat categories was compared between non-upstaged and upstaged groups (**Figure 2C, 2D, and S3)**. Using the Wilcoxon test, it was shown that there existed significant differences between non-upstaged and upstaged group when habitats considered at the duct level. The tests were carried out for both hypoxic and oxidative layers for both CA9 (**Figure 2C and 2D)**, and LAMP2b **(Figure S3**) as well as architecture, grade, lymphocytes, microcalcifications, and necrosis (**Supplementary Table 1**). As shown in Figure 2D, CA9 scoring within hypoxic habitats provides a much clearer distinction between pure DCIS and upstaged groups compared to the normoxic zone. Interestingly, CA9 did not show significant differences between the groups when analyzed at the whole duct or whole-slide level, as is traditionally done (**Figure S2B**). However, focusing on habitats revealed that CA9-positive cells are distributed differently between the two patient groups. This analysis underscores the value of examining fine-scale habitats within ducts. The improved performance of habitat scoring compared to whole-duct scoring highlights the necessity and significance of exploring the cellular composition and interactions within the tumor ecosystem.

**Figure 2.**
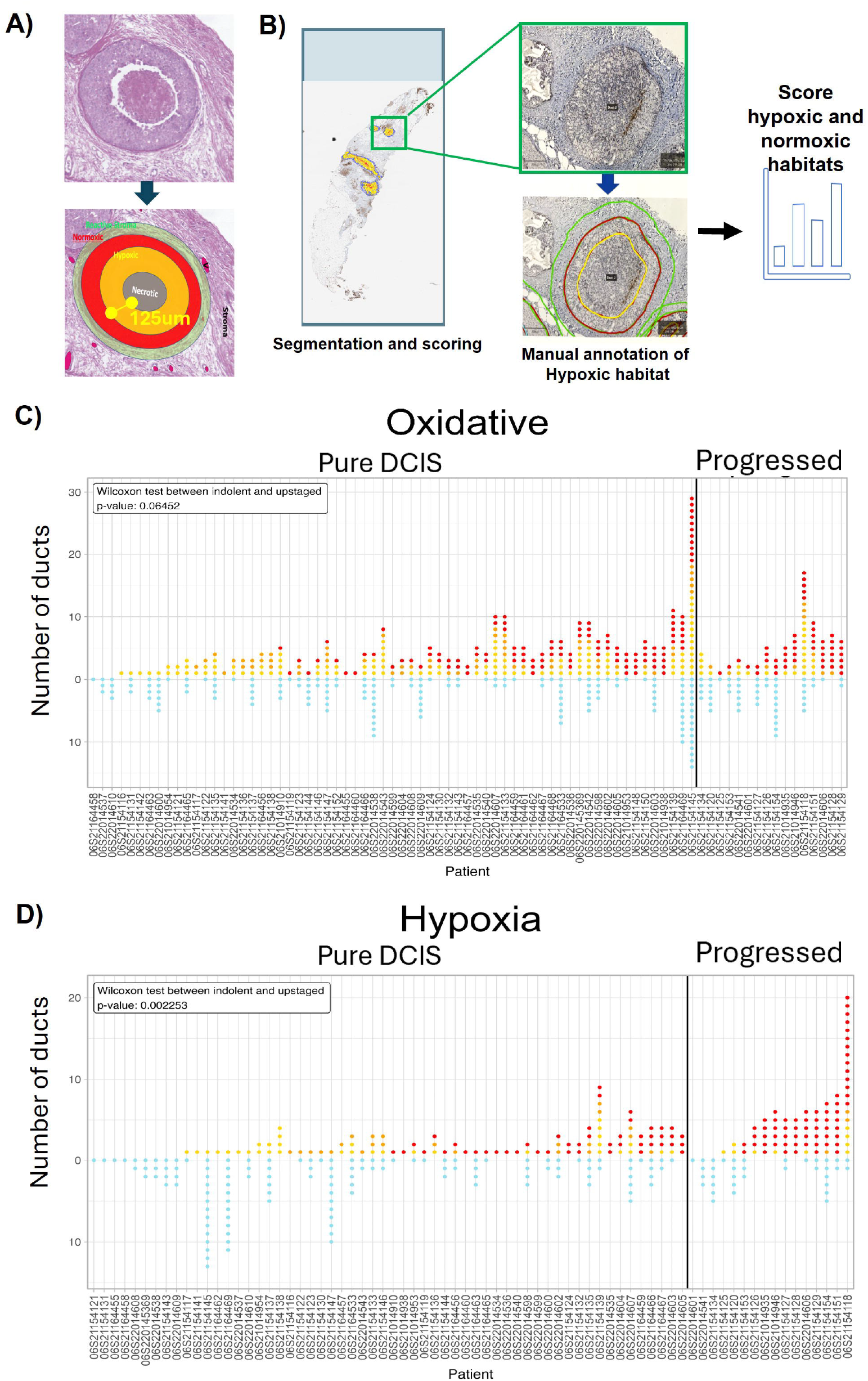
Eco-evolutionarily designed biomarker discovery to predict upstaging in DCIS. **A)** Illustration of normoxic, hypoxic and necrotic habitats in a duct. **B)** Illustration of annotation and scoring on 2 IHCs and how cells are scored in each habitat. **C)** and **D)** Dot plots of counts of CA9 expression in each habitat per duct. Cells are scored 0 for ‘negative’ or ‘1+’,’2+’,’3+’ for positive cells based on their intensity. Scoring was performed and analyzed separately for normoxic (oxidative) habitat (C) or hypoxic habitat (D). In the dot plot, each dot is a single duct. The color of dots reflects their score as follows: Blue = 0, yellow =‘1+’, orange =‘2+’, and red = ‘3+’. The number of dots reflects how many ducts were detected in each patient’s biopsy with size bigger than 400 μm in diameter. The distribution in hypoxic habitat is significantly different between pure DCIS and upstaged groups in hypoxic habitats and not in oxygenated habitats. Data was analyzed using the Wilcoxon signed-rank test. The same graph is created for LAMP2b (supplementary fig. 2).

### Mapping Metabolic Niches Within Habitats to Enhance Spatial Machine Learning Models

Previous analyses of hypoxic and normoxic habitats in ducts were limited to scoring each biomarker individually, focusing solely on the count of positive cells within each habitat. To broaden the scope and incorporate interactions and relationships between these two eco-evolutionary markers, we performed co-registration of two slides. This step enabled the creation of a virtual multiplex IHC (mIHC) by mapping cells onto a unified 2D reference space. HE slides were selected as the reference, and all IHC slides were registered onto it. (**Figure S4**). Note that since our analysis is carried out duct-by-duct, it is not necessary to register the whole slide. Instead, for each duct, we register its IHC stainings to its HE staining. This ensures all the downstream analyses could be performed on the same HE slide coordinates, providing consistency and precision in the spatial data integration. Then we used these mIHC images to define niches of cells that are positive for CA9, LAMP2b, or both. We hypothesized that niches characterized by both markers together would provide greater biological insight than analyzing each marker individually, given the established correlation between hypoxia and acid phenotypes. Then, we focus on the cell features such as nuclear morphology and texture and cell spatial features inside these niches to explore their predictive power on DCIS upstaging. As illustrated in **Figure 3**, we first map each positive cells to the reference HE slide using the co-registration described above. Then, by treating each positive cell as a node and connecting the cells within a distance threshold, we construct a cell-proximity graph out of mIHC positive cells whereby each connected component represents a continuous region or niche that is hypoxic, acidic, or both. The threshold is a tunable parameter that is optimized by the classifying power of the downstream analysis. And depending on the selection of the eco-evo markers, there can be CA9 rich niches, LAMP2b rich niches, or both CA9 and LAMP2b rich niches.

**Figure 3.**
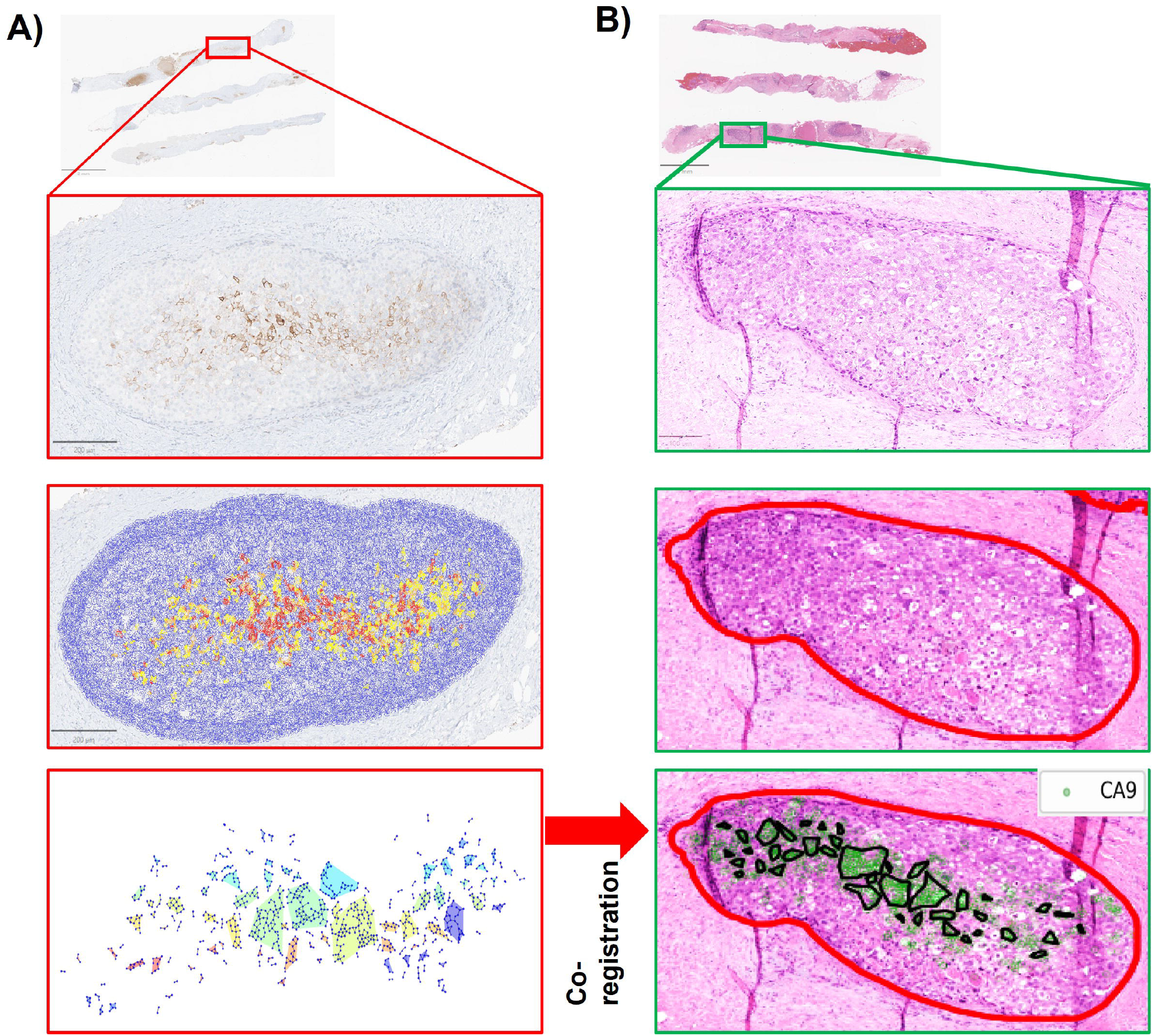
Niches are defined inside habitats from the hypoxia and acidosis markers expression. **A)** One sample duct from CA9 slide. Top: The original IHC slide. Middle: Cell detection and intensity-based classification using QuPath overlaid on the slide. Bottom: the graph constructed from the CA9 positive cells and the connected components of the graph (Niches) highlighted in different colors. **B)** The HE staining of the same duct as A. Top: The original HE slide. Middle: Duct annotation overlaid on the HE slide. Bottom: Co-registered CA9-positive niches mapped and overlaid on HE slides as mIHC to be able to extract HE features from CA9 positive niches. Note the orientation of HE and CA9 slide was opposite, and our co-registration technique successfully created a mIHC of the ducts with similar coordinates. The same approach was used for LAMP2b and the combination.

### Post Analysis reveals the top contributing patterns and features of DCIS upstaging

We then developed a pattern differential analysis pipeline, which comprises two stages: First, the samples are clustered based on the features and classified into patterns (**Figure S6**). We found eight distinct patterns shown by different color in Figure 4A. Then for each patient, we calculate the proportion of each pattern to build a distribution profile of each pattern (**figureS7**). By using these proportion features, we train a classifier to predict the upstaging (**Table 1**). We were able to predict the clinical outcome of a patient based on his/her spatially defined pattern distributions. Then, to test the hypothesis that finer regions (niche) with biological meanings could provide better predictive power, we conduct a multi scale analysis performing a series of experiments using the same set of features and with the same pattern differential analysis pipeline at 3 different scales: duct, habitat, and niche (**Figure 1C**). At the habitat level, normoxic and hypoxic zones are analyzed independently. At the niche level, analyses are further refined to separately examine CA9-positive cells, LAMP2b-positive cells, and cells positive for both. For all the experiments, the Bx dataset underwent 5-fold stratified cross-validation, where in each round, 4 folds served as the training dataset and 1-fold as the test dataset, with the goal of predicting the patients’ clinical outcome at the time of Bx. Upon comparing the mean accuracy score and the mean AUC score of all the classifiers, the niche level classifier yielded the best predictive results particularly under both metrics (**Table 1**). This result confirms that niche-based analysis outperforms our primary habitat analysis. The higher accuracy of the niche measurements may be implying that niche measurement is better than inferring habitat. Also, it is worth mentioning that oxygen habitat analysis is a rough estimate in our analysis since we do not know the exact location of the vasculature and their activity. After identifying the best-performing classifier based on the AUC metric we employed SHAP (42) (Shapley Additive exPlanations) analysis to interpret the model by calculating SHAP values for each feature, specifically on the proportions of distinct patterns. SHAP is a unified approach to interpreting machine learning models by assigning each feature an importance value for a particular prediction. In our study, SHAP values were computed for the features representing the proportions of different patterns within the niches. By calculating the SHAP values, we could determine the impact of each feature on the model’s output, thereby identifying the most influential patterns that contribute to predicting DCIS progression. This step is crucial for ensuring the transparency and interpretability of the machine learning model. Furthermore, we select features that are highly relevant to the sub-classes using different approaches including covariance, mutual information scoring and maximum relevance minimum redundancy (mRMR)(41) and choose the features identified by all three approaches. The pattern with the maximum SHAP value, identified as the most impactful, underwent further differential analysis to uncover features that significantly differentiated this pattern from others. This differential analysis employed methods including correlation analysis, mutual information (MI), and maximum relevance minimum redundancy (MRMR), which together identified F_0 <= r < 10 and solidity skewness as the top distinguishing features for Pattern 5 (**Figure 4B**). **Figure 4C** shows the gradient map of each of these features of niches in the latent space, while top differential features of other niche patterns can be found in **Figure S8**.

**Table 1.** Performance scores of multi scale classifiers. While habitat-level analysis enhanced performance, the niche-level classifier produced the most accurate predictive results.

|  | Duct | Habitat |  | Niche |  |  |
| --- | --- | --- | --- | --- | --- | --- |
|  |  | Normoxia | Hypoxia | CA9 | LAMP2b | CA9 & LAMP2b |
| **Accuracy** | $0.78 \pm 0.06$ | $0.86 \pm 0.03$ | $0.83 \pm 0.06$ | $0.82 \pm 0.06$ | $0.90 \pm 0.03$ | $0.90 \pm 0.03$ |
| **AUC** | $0.61 \pm 0.08$ | $0.67 \pm 0.03$ | $0.66 \pm 0.10$ | $0.64 \pm 0.10$ | $0.72 \pm 0.07$ | $0.74 \pm 0.13$ |

**Figure 4.**
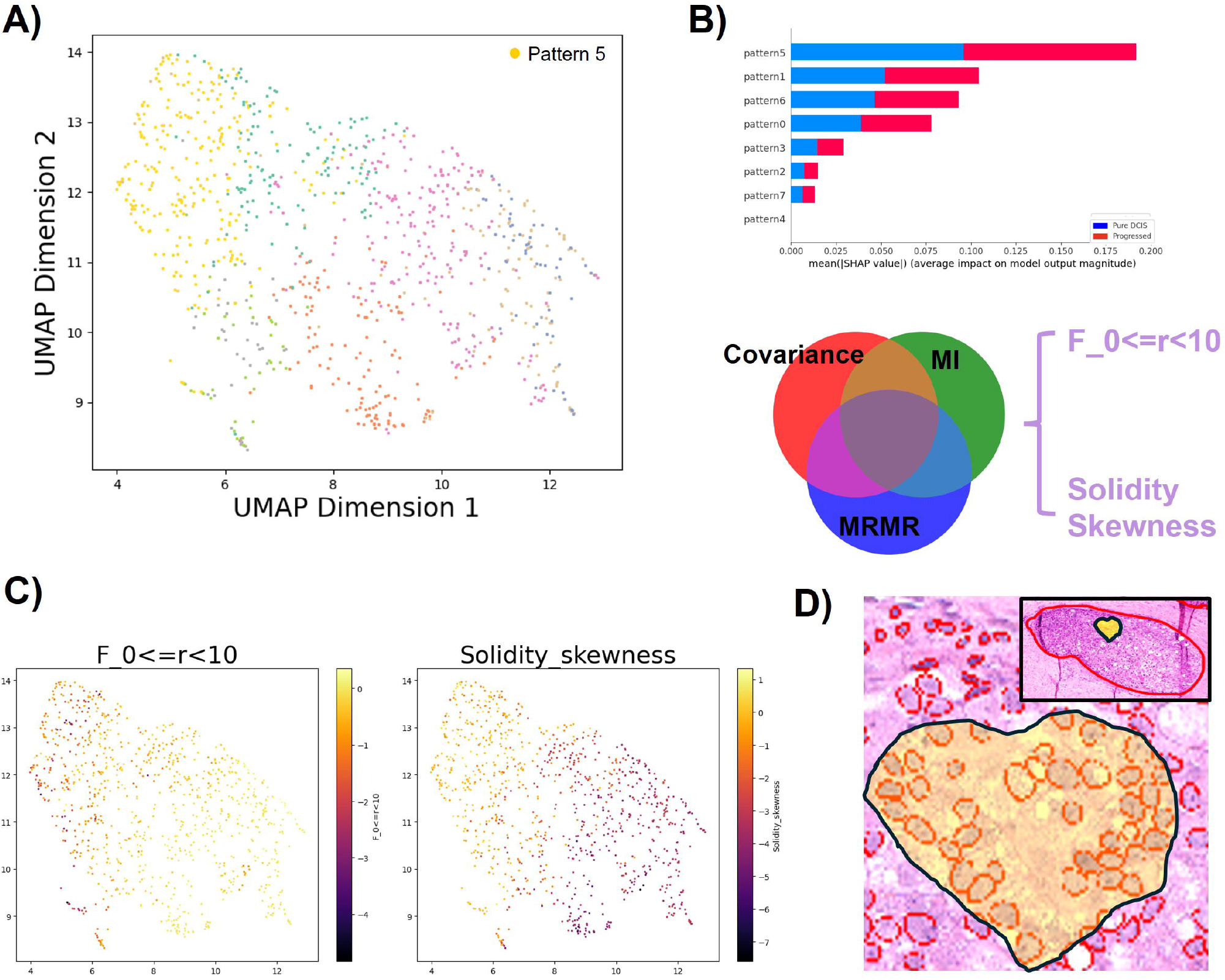
Post Analysis and projection on mIHC reveals the top contributing ecological patterns and features. **A)** UMAP (Uniform Manifold Approximation and Projection) of the features of the niches, different colors represent different patterns **B)** Top: The impact of each pattern on the classifying result, blue and red colors represent impact on pure DCIS and progressed predictions respectively, the proportion of pattern 5 has the greatest impact for both categories. Bottom: Using correlation, MI, and MRMR to obtain the most contributing features in the pattern 5 clustering phase, identifying a common feature set that includes 2 features: F_0< = r<10 and solidity skewness. **C)** UMAP showing the value of the 2 identified features for different samples, and it indicates that samples in the pattern 5 tend to have low values for F_0< = r<10 and high values in solidity_skewness. **D)** A niche belonging to pattern 5, it exhibits a relatively dispersed distribution.

### Niche distribution for diagnosis

Tumor microenvironment is heterogeneous, and niches demonstrate diverse spatial and morphological behavior. To account for the diversity, we focus on how different niches are distributed across a sample. We show that the distributions of different niches essentially characterize the tumor ecology in a much more refined manner compared with previous distance-based definitions of hypoxia/oxidative layers. After assigning each duct to its sub-class, we aggregate across all niches of each sample and use its sub-class distribution to characterize this sample. Assuming k niche sub-classes, each sample has a k dimensional histogram to describe its niche sub-class distribution. We call this the niche distributional (Nbd-Dist) feature. We trained a classifier to predict whether a sample is pure DCIS or upstaged. Repeating the iteration 10 times and comparing the mean AUC on the test set. We experimented several classifier types including lightGBM, soft vector machine (SVM), logistic regression and random forest. Random forest classifier yields the best performance on our data.

A prototype for Pattern 5, selected based on its alignment with the mean values of these features, was visualized to illustrate its characteristics (**Figure 4D**). Using a multi-scale analytical approach, we integrated spatial interactions of CA9 and LAMP2b positive cells into the machine learning pipeline to distinguish between pure DCIS and upstaged group. Niche-level analysis yielded the highest accuracy and AUC, emphasizing the importance of fine-scale regions in predicting clinical outcomes. The use of SHAP analysis and differential analysis provided an interpretable framework to highlight influential patterns and features, such as cell shape related features (Solidity) and spatial clusteredness related features (F function values), offering insights into the tumor microenvironment. This approach not only advanced our understanding of key spatial and morphological features but also demonstrated significant potential for precise diagnostic tools in clinical applications.

## Discussion

Ductal carcinoma in situ is the most prevalent type of precancer that can range from non-upstaged to aggressive. DCIS lesions are highly heterogeneous in their intra- and inter- ductal microenvironments, genetics, and molecular expression patterns. Recently, genomic analysis of matched DCIS and IDC samples has revealed that in 75% of cases, the invasive recurrence was found to be clonally related to the initial DCIS. This implies that tumor cells derived from DCIS could evolve in a linear or branching fashion with 18% new transformations and/or clonogenesis (11). These new findings emphasize the extraordinary heterogeneity in genotype and phenotypic plasticity in breast cancer that must be studied in the light of evolution and ecological studies. Tumors are complete ecosystems containing habitats and niches including normal epithelial cells, pre-cancer cells, stromal cells, vasculature, structural proteins, signaling proteins and physical factors such as pH and oxygen concentration (18). These habitats and niches can contain unique mixtures of cells with physical and biochemical characteristics, with differential evolutionary potential and trajectories (27). The niches with similar mixtures of cells usually are also similar in their physiology and phenotypes mainly due to living in similar habitats. Our hypothesis is that knowledge of these niches and their habitats can potentially provide patient benefit by stratifying their tumor progress and therapeutic choices. However, tools and techniques are lacking to distinguish them. Proper tools and techniques can identify and define habitats and niches to map precancer ecosystems.

In this study, we argue that the overdiagnosis and overtreatment of DCIS stem from conventional frameworks that focus primarily on genetic signatures while neglecting the ecological heterogeneity within tumor ecosystems. Thus, we interpreted complex eco-evolutionary data of cancer cells within their niche using machine learning and pathomics, all framed within an innovative ecological and evolutionary dynamic model. Oxygen habitats are identified based on varying levels of perfusion and oxygenation, which are believed to play a crucial role in driving ecological diversity by changing cancer cells metabolism, creating new habitats, and enhancing tumor heterogeneity, ultimately leading to diverse evolutionary trajectories (28, 29). Solid tumors often exhibit an impaired vascular system, leading to habitats within tumors that vary in hypoxia, nutrient deficiency, and acidity. These habitats can significantly influence the spatial selection of cellular phenotypes in distinct subregions. Inhabiting hypoxia, acidosis, and severely nutrient deprived, face (pre-)cancer cells to strong selective pressures leading to divergence to novel phenotypes in population. These new phenotypes can reciprocally influence the microenvironment resulting in a dynamically changing tumor ecosystem. Therefore, the phenotype of the cells residing in these habitats can also be leveraged to define the habitats with a certain degree of accuracy. Previous research from our group and others demonstrated that cancer cells within breast ducts, exposed to chronic hypoxia and acidosis, develop adaptive mechanisms for survival in this challenging microenvironment including expression of CA9 or LAMP2b at the cell surface (18,20,30). However, none of these findings were used in a relevant translational study for biomarker discovery. In this study, we explore these biomarkers within an eco-evolutionary framework for the first time, using them as indicators of the metabolic state of cancer cells residing in a niche as part of oxygen habitats that may favor the selection of more aggressive phenotypes to predict the upstaging of DCIS. While a longitudinal study would indeed be a better study design for direct observation of evolutionary changes over time, our current cross-sectional approach enables us to capture a snapshot of the tumor microenvironment at two near time points, providing valuable insight into the conditions that distinguish DCIS from IDC. We recognize the assumption that the synchronous IDC microenvironment may contribute to the progression from DCIS to IDC. However, our study design allows us to test whether specific microenvironmental factors and related habitats and niche correlate with the presence of IDC, which can provide strong hypotheses for future longitudinal investigations. A future prospective or retrospective longitudinal (multiple long time points) study would indeed help distinguish whether these microenvironmental changes in tumor ecosystem locally belonged to habitats or niches can drive progression from DCIS to IDC or if IDC-induced those changes in the tumor ecosystem contribute to the synchronous DCIS phenotype.

In our curated retrospective cohort of 84 DCIS patients with histologically confirmed DCIS on core biopsy, we manually annotated 916 single ducts and more than 3000 habitats on all three slides and scored them at habitat levels. This unique detailed eco-evolutionary annotation can be used for future similar eco-evolutionary designed studies including reactive stroma habitats. Our risk scoring system integrating principles of ecological-evolutionary dynamics with pathological imaging and molecular features of early-stage breast tumors showed improvement on prediction power of biomarkers alone and in combination.

We employed a 5-fold stratified cross-validation approach to ensure robust internal validation of our model. While this method helps mitigate overfitting and provides reliable performance estimates, we acknowledge the absence of an independent validation set, which is crucial for assessing the model’s generalizability. The unique design of our cohort, which integrates specific ecological and microenvironmental factors, limits the availability of comparable external datasets for validation. As such, there is no current dataset with similar characteristics for cross-validation. We recognize this as a key limitation and emphasize that future studies should aim to validate the model on independent cohorts when such datasets become available. Furthermore, although our model achieved an AUC of 0.74, this performance is not yet sufficient for clinical translation. Additional efforts to refine the model and test it in larger, independent cohorts will be essential before its use in clinical practice can be considered. Interestingly, a recent approach using multiplex IF on DCIS cohort reached the same AUC(2). While both our study and the Risom et al. paper aim to leverage spatial relationships to predict DCIS progression, we would like to emphasize that the two approaches are fundamentally different in terms of the markers used. Risom et al. focused on a broad panel of markers, including those related to the stroma, immune cells, and tumor cells, which provide a comprehensive view of the tumor microenvironment. In contrast, our approach centers on eco-evolutionary markers derived from adaptation of cancer cells to physical microenvironment, specifically CA9 and LAMP2b, which are associated with hypoxia and tumor acidity and their spatial distribution, respectively. These differences reflect divergent hypotheses about the key drivers of DCIS progression. The fact that both studies report a similar AUC of 0.74, with the distinct marker sets and biological processes, suggests that our findings offer complementary insights into DCIS progression and combination of approaches might increase the accuracy.

Our study demonstrates the utility of eco-evolutionary principles in understanding DCIS progression. In our study, we proposed that specific tumor microenvironmental conditions, such as hypoxia and acidosis, are associated with phenotypic changes that may indicate DCIS progression. However, although we have shown previously that these microenvironments can cause aggressive phenotypes, we acknowledge that our findings here do not conclusively demonstrate that these environmental factors are causative agents in the transition from DCIS to IDC. Instead, our data suggests that these conditions could serve as biomarkers for identifying lesions that are more likely to be upstaged. However, the ability to define more refined cell phenotypes within each region of interest (ROI) could further enhance our analysis. If we can identify and characterize more detailed phenotypes, it would allow us to extract additional features that describe the spatial interactions of these phenotypes. This, in turn, could potentially improve the classifier’s performance and make the results more interpretable. By capturing the intricate interactions between various cell types and their microenvironments, we could gain deeper insights into the ecological dynamics driving DCIS progression and improve predictive models for patient outcomes.

In recent years, there has been a growing trend towards adopting a “watchful waiting” approach for certain cases of DCIS, rather than immediate surgical excision (31,32). This strategy aims to reduce overtreatment by closely monitoring DCIS lesions that may not progress to invasive cancer. In this context, our upstaging predictions become particularly relevant. Identifying microenvironmental and phenotypic factors that indicate a higher likelihood of progression to IDC could help clinicians make more informed decisions about when to intervene and when to adopt a more conservative, observational approach. The ability to predict which DCIS cases are at higher risk of progressing to invasive disease would provide critical information for optimizing patient management, minimizing unnecessary treatments, and reducing the psychological and physical burdens associated with overtreatment (33). Further validation of these predictive models could therefore have important implications for guiding treatment strategies in the context of DCIS.

### Lead contact

Further information and any related requests should be directed to and will be fulfilled by the lead contact Mehdi Damaghi.

## Supporting information

All supplementary images

## Acknowledgements

We gratefully acknowledge funding from Physical Sciences Oncology Network at the National Cancer Institute (grant U01CA261841), R01 grant R01CA272601, and R01CA249016. This work has been supported also in part by the Analytic Microscopy Core Facility at the H. Lee Moffitt Cancer Center & Research Institute, an NCI designated Comprehensive Cancer Center (P30-CA076292). We thank Dr. Alexander Borowsky and Dr. Kenneth Shroyer, for reading our manuscript and constructive comments on pathological aspect as well as Dr. Liliana Davolos and Dr. David Ray for their guidelines on ecological and evolutionary principles used in the manuscript.

## Author contributions

M.D. conceptualized and designed the research; Y.X., M.A., J.D.B., A.C., Y.C., M.D., performed the experiment and analysis; J.D.B. reviewed all the slides and scored them as the project pathologist; P.P., C.C., and M.D. contributed to results interpretation; and M.D. wrote the paper. All authors revised the paper.

