## Supplementary material for "Eco-evolutionary Guided Pathomics Analysis to Predict DCIS Upstaging": All supplementary images

**Supplementary information:**

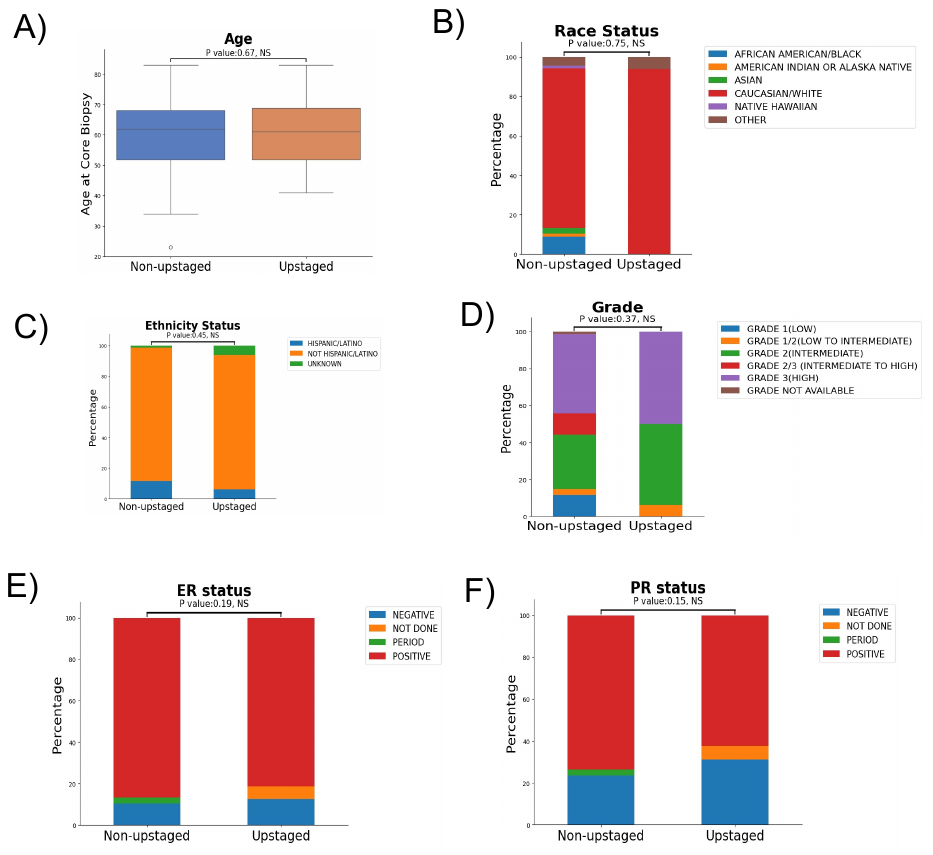

**Figure S1. Clinical features comparison between non-upstaged and upstaged groups.** A) Boxplot of patients’ ages for ‘non-upstaged’ group and ‘Upstaged’ group, p-value of Wilcoxen test is 0.6783. B) Race status, p-value: 0.7626; C) Ethnicity status, p-value: 0.4982; D) Grade, p-value: 0.3777; E) ER status, p-value: 0.2126 F) PR status, p-value: 0.1775. B-F) Stacked bar plot of clinical features of the two groups is shown and chi-square tests are performed for analysis.

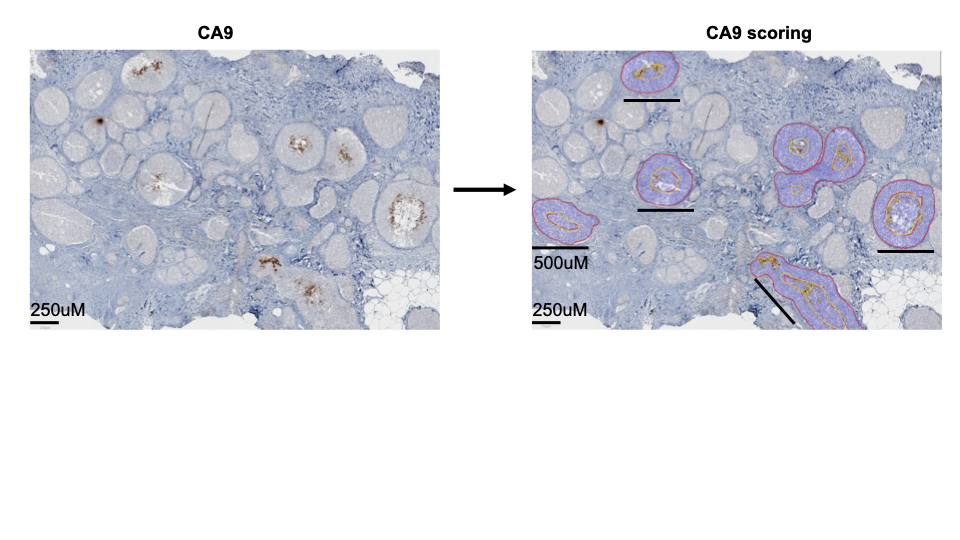

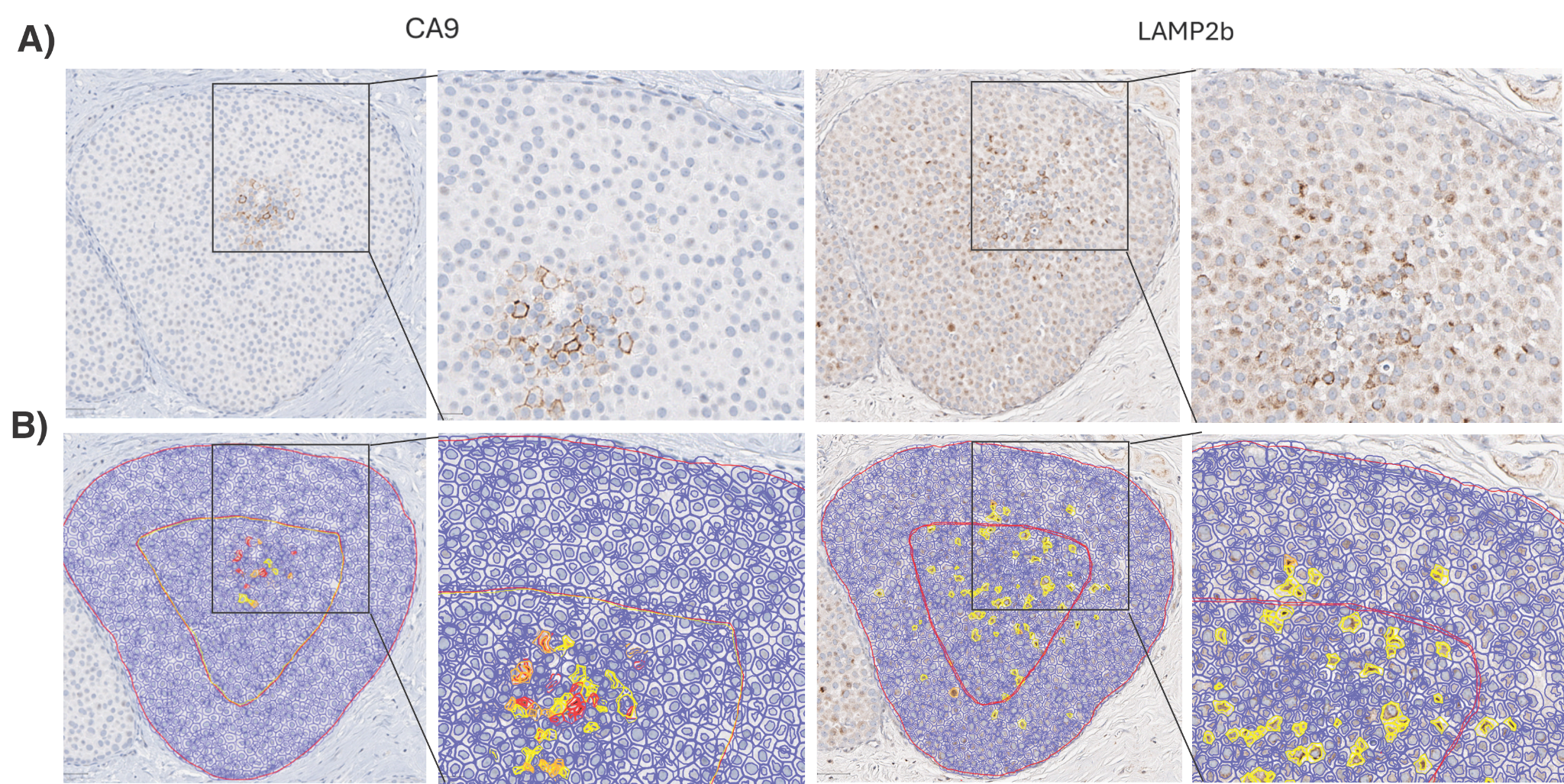

**Figure S2.** **Duct manual annotation and scoring.** Ducts bigger than 400 uM in diameter were manually annotated in two zones of normoxia and hypoxia representing oxygenated and hypoxic habitats. Blue is CA9 and LAMP2b negative and yellow, orange, and red are +1, +2, and +3 positive respectively.

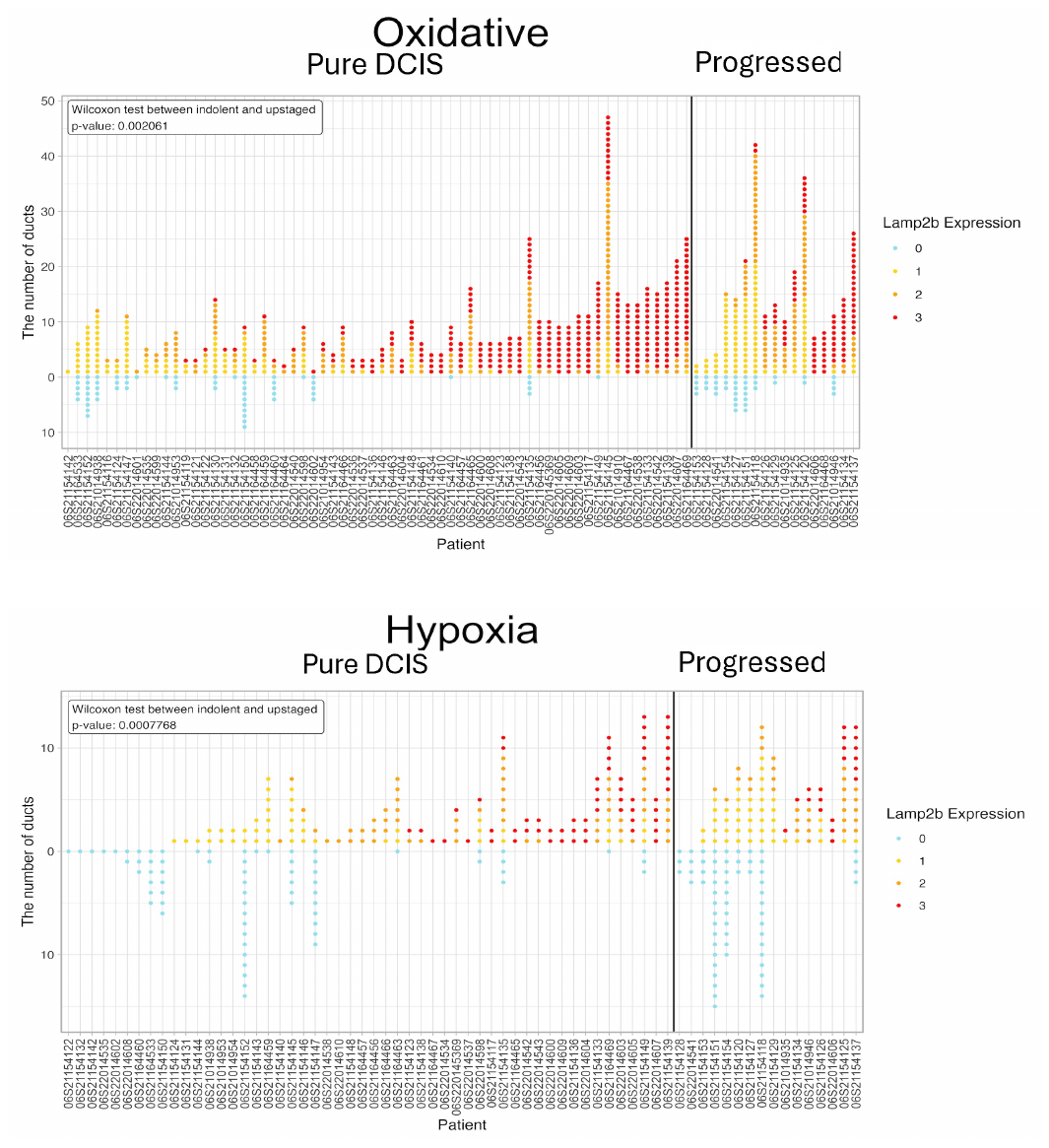

**Figure S3. LAMP2b scored duct counts.** Dot plots of counts of LAMP2b ‘negative’,’1+’,’2+’,’3+’ scored ducts for all the biopsy samples. scoring was done on normoxic habitat(top) and hypoxic habitat (bottom).

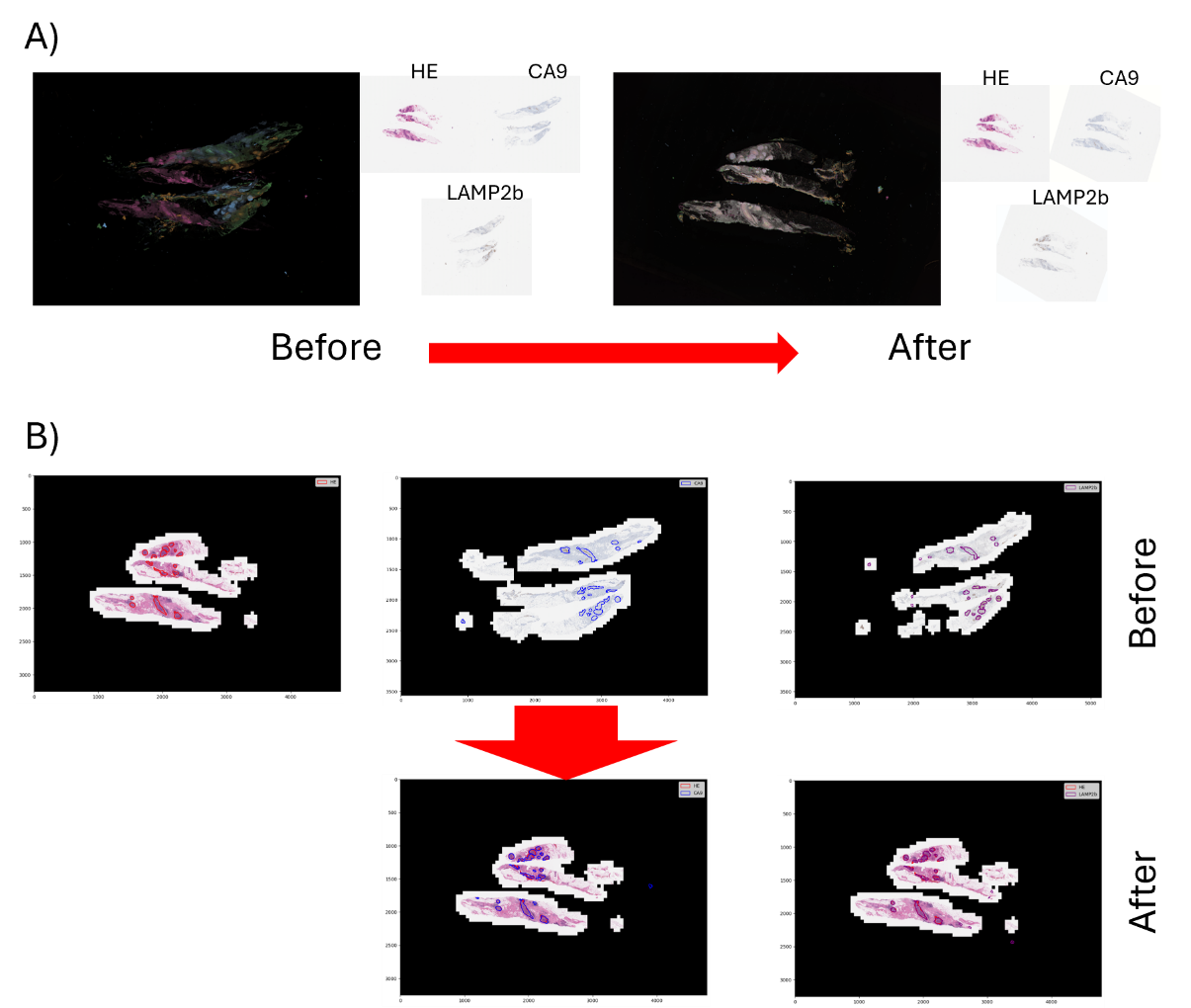

**Figure S4.** **Co-registration of HE with IHC slides.** A) Slides before and after slide-level (VALIS) co-registration. Left: Original slides, right: rotated and warped slides. It shows clearly the slide level co-registration aligns the images’ orientations and the tissue locations B) Duct annotations before and after duct-level co-registration. Top: original annotations on HE and IHC slides, bottom: co-registered annotations on HE slides.

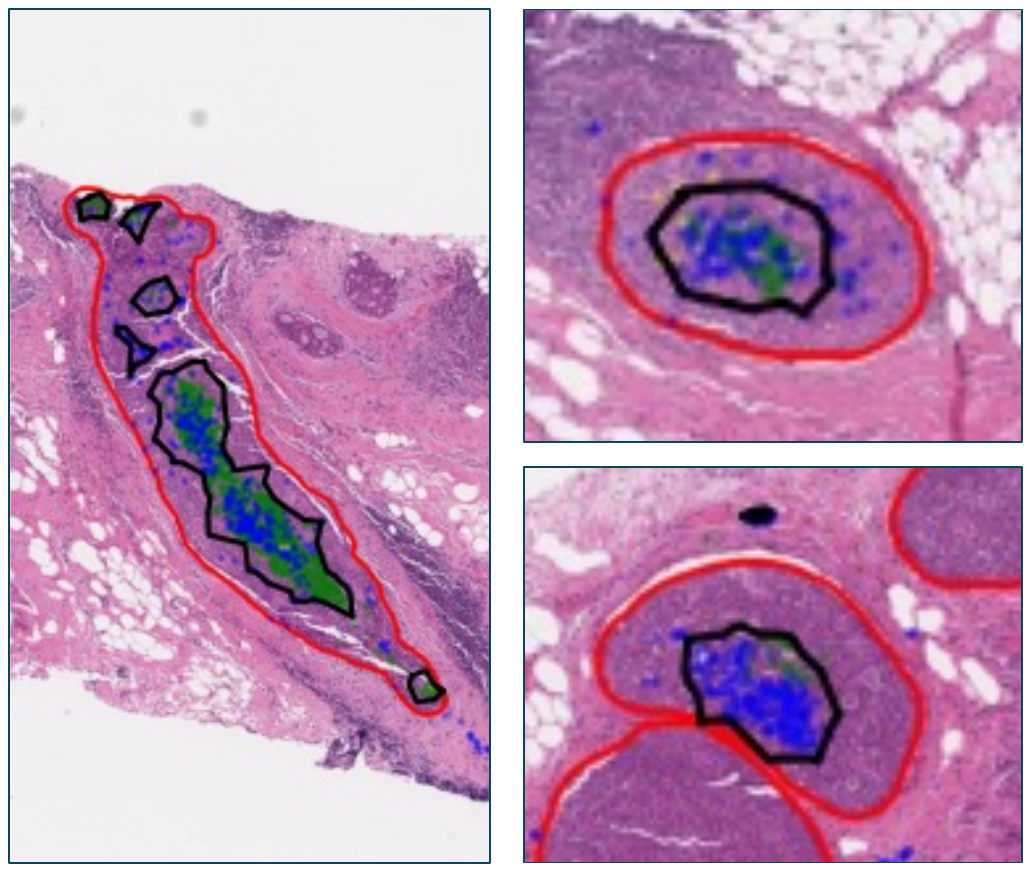

**Figure S5. Visualization of niches overlaid on HE image.** Co-registered CA9 positive cells and LAMP2b positive cells are overlaid on HE slide, represented by green and blue dots, constructed niches are outlined in black.

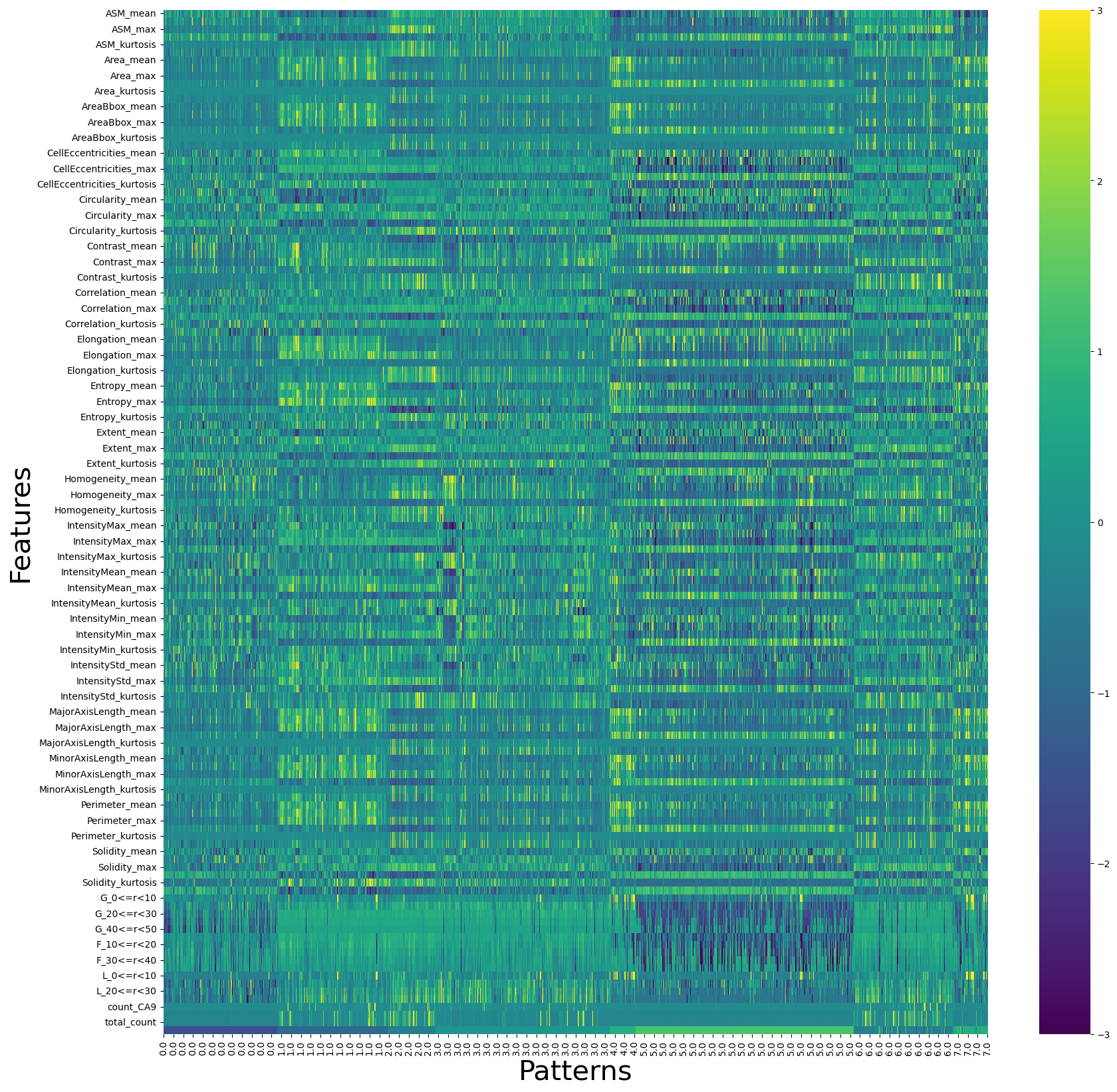

**Figure S6. Heatmap of Feature Values Across Different Niche Patterns.** The heatmap displays the values of features for each niche (sample), with rows representing features and columns representing niches. The x-axis ticks indicate the pattern to which each niche belongs. The color intensity in the heatmap corresponds to the feature values, illustrating the variation in feature expression across niches and patterns.

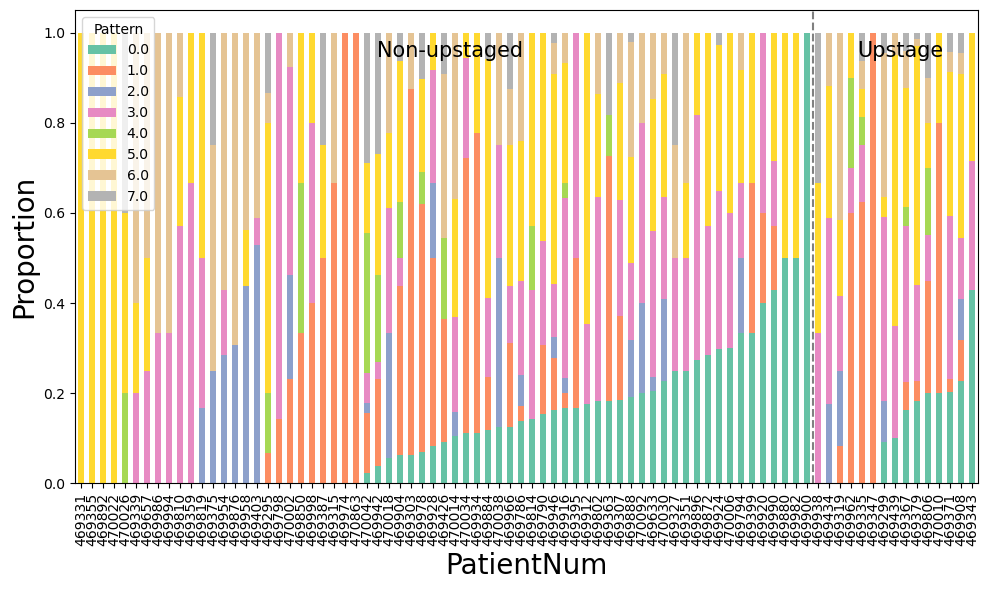

**Figure S7. Distribution of Niche Patterns by Patient Group: Non-Upstaged vs. Upstaged.** The stacked histogram displays the proportion of niches associated with different patterns for each patient. Each bar represents a patient, with the stacked segments indicating the proportion of niches for each pattern. Patients are grouped into Non-Upstaged (left) and Upstaged (right), displaying a comparison of pattern distributions between the two groups.

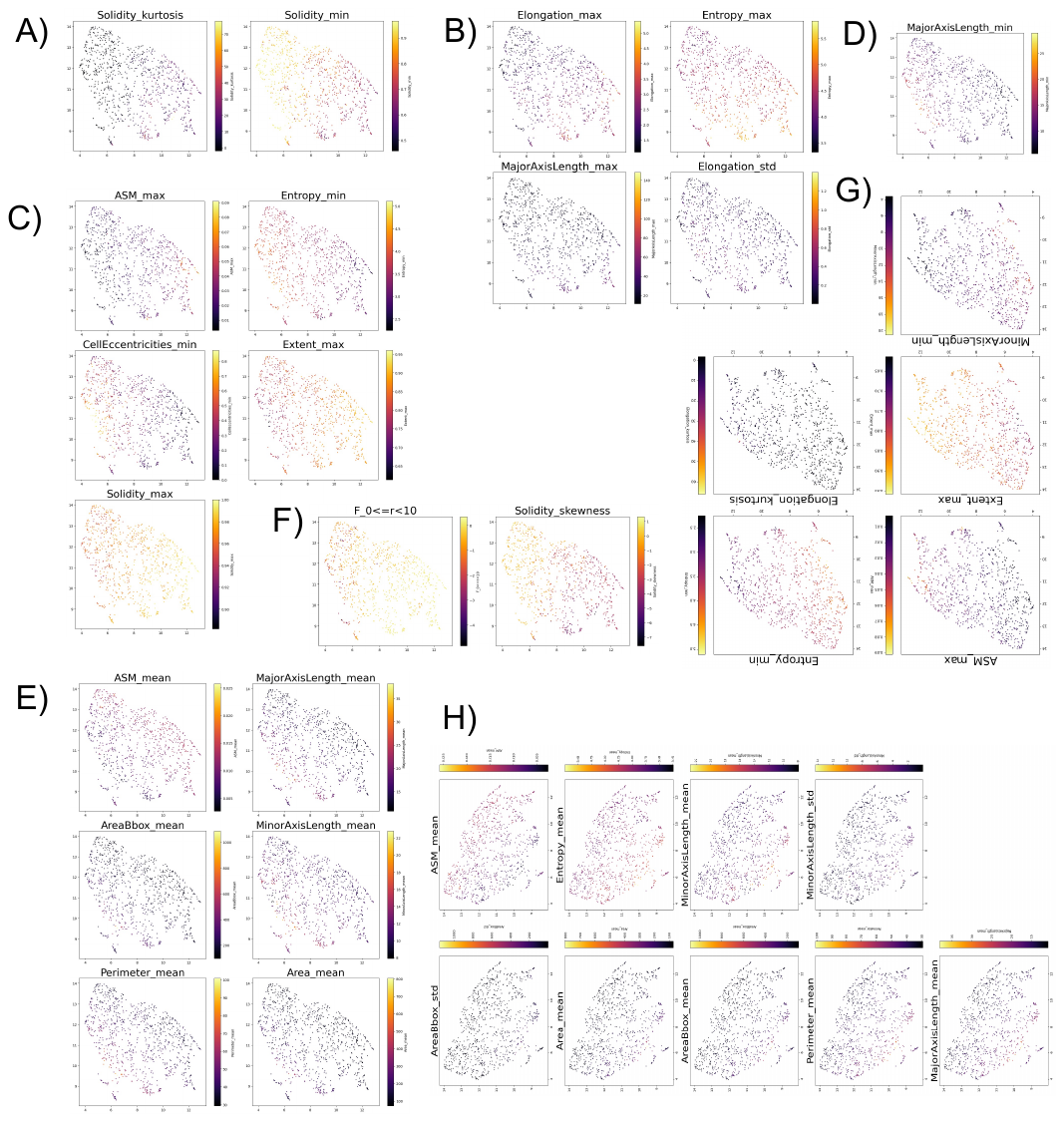

**Figure S8**. **Gradient Maps of Top Differentiating Features for All Patterns.** (A) to (H) represent Pattern 0 to Pattern 7, respectively. Each subfigure shows a UMAP visualization colored by the values of the most differentially expressed features for the corresponding pattern.
